# STAT3 and HIF1α cooperatively mediate the transcriptional and physiological responses to hypoxia

**DOI:** 10.1101/2021.11.19.469257

**Authors:** Alberto Dinarello, Riccardo Massimiliano Betto, Linda Diamante, Chiara Cioccarelli, Giacomo Meneghetti, Margherita Peron, Annachiara Tesoriere, Claudio Laquatra, Natascia Tiso, Graziano Martello, Francesco Argenton

## Abstract

STAT3 and HIF1α are two fundamental transcription factors involved in many merging processes, like angiogenesis, metabolism, and cell differentiation. Notably, under pathological conditions, the two factors have been shown to interact genetically, but both the molecular mechanisms underlying such interactions and their relevance under physiological conditions remain unclear. Here we report that STAT3 is required for the HIF1α-dependent response to hypoxia. In Stat3 knock-out pluripotent embryonic stem cells (ESCs), a large fraction of HIF1α target genes is not induced by hypoxia. Mechanistically, STAT3 does not regulate neither HIF1α expression nor stability, rather, it physically interacts with it in the nucleus. *In vivo*, we observed that both genetic and chemical inactivation of Stat3 blunted physiological responses to hypoxia, such as angiogenesis, erythropoiesis, and immune cell mobilization. Such defects were accompanied with faulty transcriptional activity of HIF1α. In sum, our data reveal that STAT3 and HIF1α cooperatively mediate the physiological response to hypoxia.

## INTRODUCTION

Hypoxia Inducible Factor (HIF) 1α, the master regulator of cell metabolism in response to low levels of available oxygen, is involved in a wide range of biological processes, such as cancer progression, angiogenesis, and differentiation of pluripotent stem cells (Semenza, 2012; Zhou *et al*., 2012). HIF1α is constitutively expressed and generally degraded by Von Hippel Lindau tumor suppressor protein (pVHL) upon hydroxylation by Prolyl Hydroxylase Domain-containing (PHD) enzymes (Masson and Ractliffe, 2003; Kaelin, 2017). In hypoxic conditions PHD3 is unable to hydroxylate HIF1α that is therefore stable enough to migrate in the nucleus where it generates heterodimers with HIF1β, a stable and constitutively expressed protein (Semenza, 2001). The HIF1α/HIF1β heterodimer activates the transcription of target genes involved in angiogenesis, cell metabolism, differentiation of pluripotent stem cells, erythropoiesis, and immune cell migration (Rey and Semenza, 2010; Katsuda *et al.,* 2013; Pimpton *et al*., 2015; Haase, 2013; Gerri *et al*., 2017; Zhuo *et al*., 2012). In the last decade, *in vitro* studies revealed connections between HIF1α and other transcription factors among which Signal Transducer and Activator of Transcription 3 (STAT3) emerged as an important modulator of hypoxia/HIF1α activity (Pawlus *et al*., 2014; Grillo *et al*., 2020). STAT3 is a component of the JAK/STAT pathway that has been claimed to control cell proliferation in cancer and pluripotent stem cells, *via* their transcriptional and metabolic regulation (Avalle *et al*., 2017; Tengesdal *et al*., 2021; Carbognin *et al*., 2016; Betto *et al*., 2021, Tesoriere *et al.,* 2021).

Under pathological conditions, the crosstalk between JAK/STAT3 and hypoxia/HIF1α pathways is significant: rheumatoid arthritis is an autoimmune disease characterized by high levels of inflammation due to the crosstalk between STAT3 and hypoxia pathways (Gao *et al.,* 2015); glucose deprivation in brain pericytes is regulated by STAT3-mediated induction of HIF1α (Carlsson *et al*., 2018); the self-renewal of glioma stem-like cells, which was previously considered as a hypoxia-dependent mechanism, appeared to be determined by an upregulation of STAT3 by HIF1α (Almiron Bonnin *et al*., 2018). Notably, targeting of STAT3 cascade can block HIF1 and VEGF signaling induced by several oncogenic pathways (Xu *et al*., 2005).

Additionally, hypoxia has a role in differentiation and proliferation of mouse embryonic stem cells (ESCs): in particular, short exposures to low oxygen tension determine the differentiation of stem cells into definitive endodermal cells and distal lung cells (Pimton *et al*., 2015); it has been demonstrated that hypoxia and retinoic acid can induce differentiation of mouse ESCs in hepatic progenitor cells (Katsuda *et al*., 2013); furthermore, HIF1α has been shown to drive the developmental progression of mouse ESCs (Zhou *et al*., 2012). Mouse ESCs are characterized by high levels of expression of *Stat3*, which is involved in the maintenance of pluripotency (Matsuda *et al*., 1999). Indeed, mouse ESCs are cultured in presence of Leukemia Inhibitory Factor (LIF), a cytokine that activates the JAK/STAT3 pathway increasing ESCs self-renewal (Ying *et al.,* 2008; Martello *et al.,* 2013; Wray *et al.,* 2010; Dunn *et al.,* 2014). Mouse ESCs can also be maintained in a medium containing GSK3 and MEK inhibitors (2i conditions). The use of 2i allowed for the derivation and expansion of *Stat3* knock-out ESCs (Ying *et al*., 2008), revealing a role of STAT3 also in the control of metabolism and proliferation (Carbognin et al., 2014; Betto *et al.,* 2021; Huang *et al*., 2014).

Despite several studies indicating a functional interplay between HIF1α and STAT3 under pathological conditions, it is not clear whether and how the two transcription factors regulate each other. For this reason, we decided to study at the molecular level the crosstalk between HIF1α and STAT3 in the context of the physiological response to hypoxia, taking advantage of genetic models developed in ESCs and in zebrafish.

## RESULTS

### STAT3 regulates the expression of a specific subset of hypoxia-dependent genes

To analyze the involvement of STAT3 in hypoxia-dependent processes, we decided to perform RNAseq and compare the data of *Stat3^+/+^* and *Stat3^-/-^* cells incubated for 24 hours either in normoxia or in hypoxia (1% O_2_ tension). As shown in Fig. 1A and in Fig. S2, the comparison between *Stat3^+/+^* ESCs grown in normoxia or hypoxia revealed a relevant group of differentially expressed genes. In particular, as reported in the Volcano plot (Fig. 1A), the expression of about 1’500 genes is altered in hypoxia and a large part of these genes are upregulated. As expected, genes involved in glycolysis and angiogenesis belong to the subset of upregulated genes. Among them we found *Vascular Epithelial Growth Factor A* (*Vegfa*), *Hexokinase* 1 (*Hk1*), *Hexokinase 2* (*Hk2*), *Phosphofructokinase* (*Pfkp*), *Lactate dehydrogenase A* (*Ldha*)*, Aldolase A* (*Aldoa*), *Phosphoinositide-dependent kinase 1* (*Pdk1*) (Fig. 1A). Interestingly, when comparing the number of genes affected by hypoxia in *Stat3^+/+^* and in *Stat3^-/-^* ESCs, we saw that the effect of hypoxia in *Stat3* mutant cells was significantly attenuated: as reported in the box plots of Fig. 1B and B’, only 70% of hypoxia-dependent genes are induced by hypoxia in *Stat3* null cells (FC>1 and p-value<0.05). Hence, 30% of hypoxia-responsive genes are not significantly induced when *Stat3* is knocked-out. Of note, the comparison between hypoxic and normoxic *Stat3^-/-^* cells revealed a significant difference in both down- and up-regulated transcripts, however, this response to low oxygen was weaker when compared to the one observed in hypoxic *Stat3^+/+^* ESCs (Fig. 1B,B’). Indeed, the comparison between hypoxic S*tat3^+/+^* and hypoxic *Stat3^-/-^* cells allowed us to identify a subset of hypoxia responsive genes whose induction is significantly dampened in *Stat3^-/-^* cells. This subset of genes is reported in the heatmap of Fig. 1C and contains canonical Hif1α targets, such as *Vegfa*, *Hk1*, *Hk2*, *Pfkp* and *Hypoxia lipid droplet-associated* (*Hilpda*) whose expressions were plotted in Fig. 1D and validated with RT-qPCR in Fig. 1E. Moreover, as shown in Fig. S1, the comparison between *Stat3^+/+^* and *Stat3^-/-^* hypoxic ESCs shows that hypoxia does not affect the expression of STAT3 target genes: as reported in Fig. S1, volcano plot and bar plots reveal that the number of STAT3-dependent genes does not change when cells are incubated in normoxia or in hypoxia. Box-plots reported in Fig. S1B demonstrated that hypoxia does not significantly affect the expression of STAT3-related genes. *Stat3* itself and some of its target genes, such as *Socs3* and *Klf4,* are downregulated in *Stat3* knock-out cells, but they are not differentially expressed when comparing normoxic and hypoxic *Stat3^+/+^* cells (Fig. S1D,E,F). Only *Tet2* is significantly downregulated by hypoxia in *Stat3^+/+^* ESCs, but this result is not confirmed by RT-qPCR (Fig. S1G). We can conclude that STAT3 is plays an important role in the induction of hypoxia-dependent genes, including known HIF1α-targets involved in the regulation of glycolytic metabolism and in vascular remodeling (Fig. S2B), while HIF1α does not regulate STAT3 activity in mouse ESCs. It is worth mentioning that some HIF1α-dependent genes are affected by hypoxia either in *Stat3^+/+^* and *Stat3^-/-^* cells. In Fig. S2C we show that the expression of *Egln3*, one of the main targets of HIF1α which encodes for PHD determining the elastic feedback loop of the oxygen deprivation response (Minamishima *et al*., 2009; Walmsley *et al*., 2011; Santhakumar *et al*., 2012), is not affected by *Stat3* mutation.

**Fig. 1:**
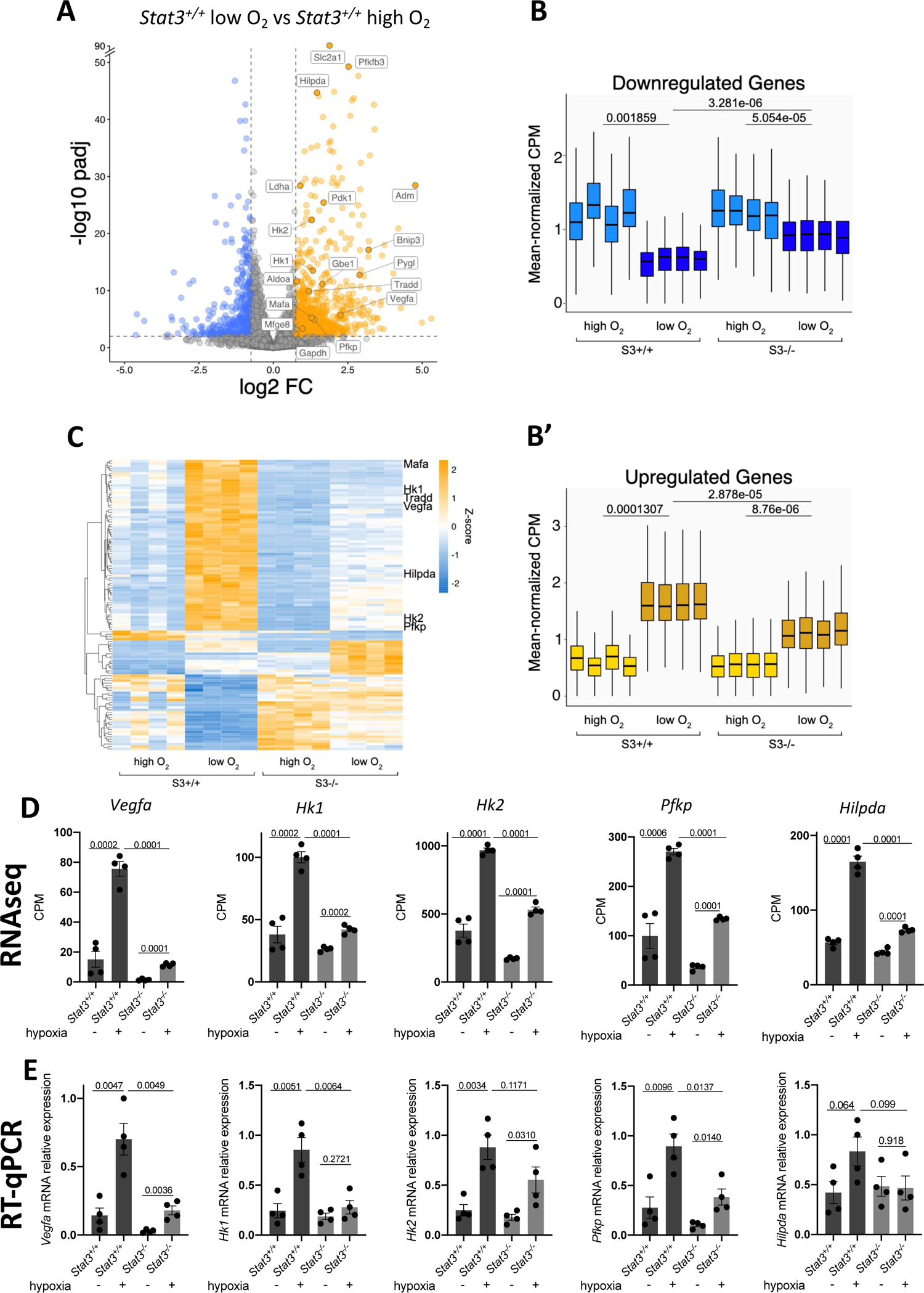
STAT3 regulates a subset of hypoxia-dependent genes. A–D: Transcriptome analysis by RNA-seq. A: Genes that were differentially expressed (log2[fold change (FC)] > + 0.75 or < −0.75; q-value < 0.01, Benjamini–Hochberg adjustment, as indicated by dashed lines) between *Stat3^+/+^* cells in normoxia and Stat3*^+/+^* cells in hypoxia; n = 4 biological replicates. B: Expression levels of genes that were down-(B) and upregulated (B’) in *Stat3^+/+^* in hypoxia relative to *Stat3^+/+^* cells in normoxia. Each boxplot shows the 1st, 2nd and 3rd quartiles; the whiskers show the minimum and maximum values. C: heatmap of RNAseq data reporting the expression of genes that are affected by hypoxia in *Stat3^+/+^* cells compared to *Stat3^+/+^* normoxic cells, but that require STAT3 to be properly altered in their expression. D: expression level of *Vegfa, Hk1, Hk2, Pfkp* and *Hilpda* in normoxic and hypoxic *Stat3^+/+^* and *Stat3^-/-^* mESCs taken from RNAseq data. E: Gene expression analysis by RT–qPCR of *Stat3^+/+^* and *Stat3^-/-^* cells treated in normoxic and hypoxic mESCs of the same genes shown for the RNAseq. Mean±SEM of n = 4 experiments, with each replica shown as a dot. *p<0.05, **p<0.01, ***p<0.001, ****p<0.0001.

To understand how STAT3 regulates hypoxia-dependent gene expression, we decided to study the crosstalk between STAT3 and HIF1α. We first asked whether the expression levels of *Hif1α* mRNA are affected by the genetic ablation of *Stat3*. Interestingly, as reported in Fig. 2A, there are no significant differences in the expression of *Hif1α* in *Stat3^-/-^* cells compared to *Stat3^+/+^*. Therefore, we sought to assess whether STAT3 is involved in the stabilization of HIF1α in hypoxia. Interestingly, as reported in Fig. 2B, western blot analysis revealed that hypoxia can stabilize HIF1α both in *Stat3^+/+^* and in *Stat3^-/-^* cells. Once demonstrated that in mouse ESCs STAT3 is neither involved in the expression of *Hif1α* mRNA nor in HIF1α protein stabilization, we decided to test whether STAT3 might interact directly with HIF1α. To do so, we performed Proximity Ligation Assay (PLA) and found that PLA positive dots could be detected only in murine *Stat3^+/+^* ESCs grown at low oxygen tensions (Fig. 2C), revealing that STAT3 and HIF1α interact with each other in hypoxic conditions. Furthermore, the cellular localization of dots indicates that the interaction between the two transcription factors occurs in the nucleus.

**Fig. 2:**
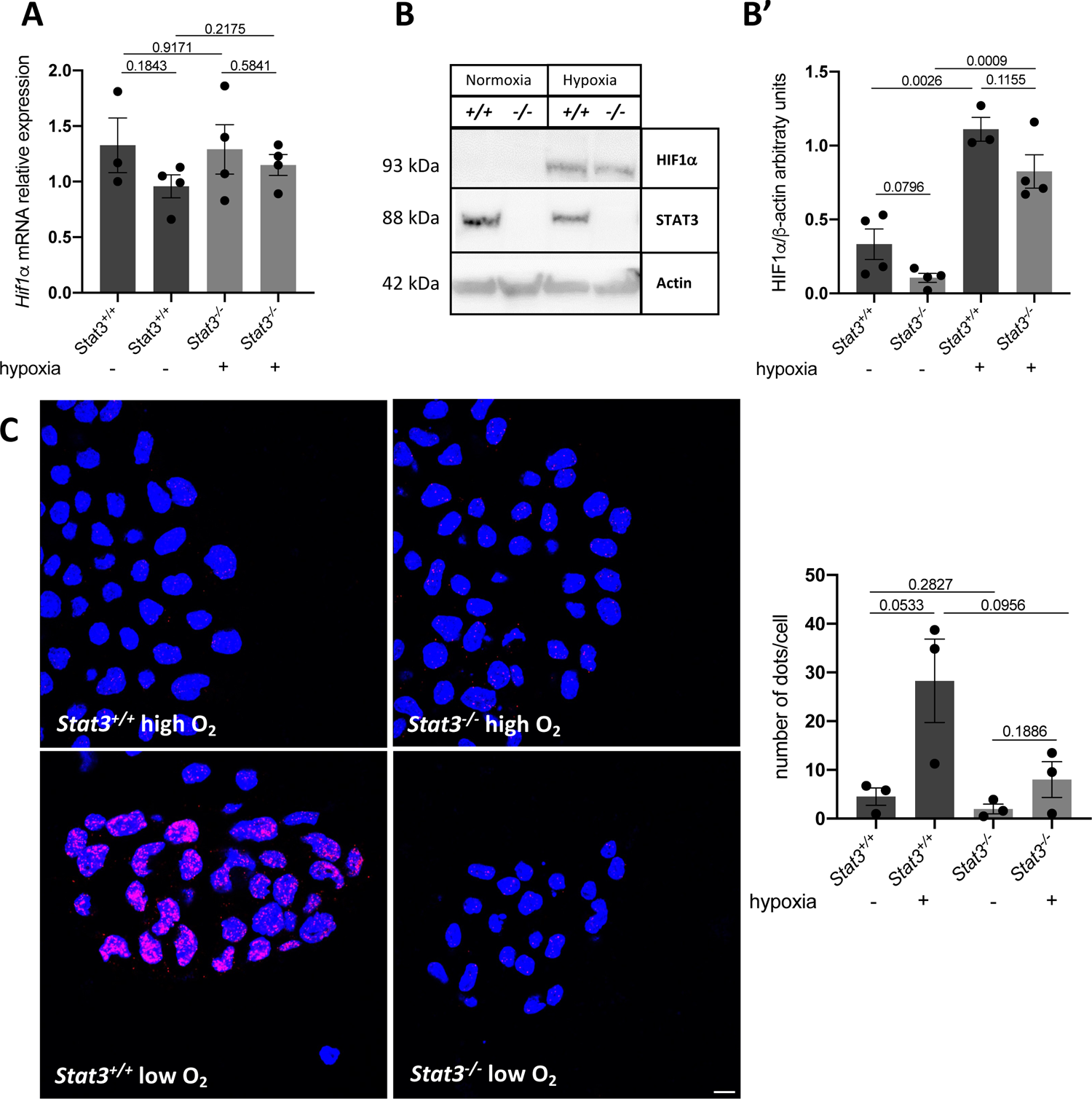
STAT3 physically interacts with HIF1α. A: RT-qPCR analysis of *Hif1α* mRNA expression on murine *Stat3^+/+^* and *Stat3^-/-^* murine ESCs incubated either in low and in high oxygen tensions. B: western blot analysis of HIF1α and STAT3 from protein extracts of *Stat3^+/+^* and *Stat3^-/-^* murine ESCs incubated either in low and in high oxygen tensions. Representative pictures (B) and quantifications (B’). β-Actin was used as an internal control. 4 independent biological replicates were used. C: PLA with anti-STAT3 and anti-HIF1α antibodies (scale bar=200 μm). Quantification of dots divided for the number of nuclei detected with DAPI (blue) (C’). n=3 independent biological replicas were used. Scale bar = 200 μm. Mean±SEM. *p<0.05, ***p<0.001; ns = not significant.

Next, we tested whether the interaction between STAT3 and HIF1α can be observed also in pseudohypoxia. Pseudohypoxia is a condition that occurs whenever HIF1α is stabilized even if the organism is exposed to atmospheric oxygen tensions. This phenomenon can be chemically triggered by some compounds like cobalt chloride (CoCl_2_) which substitutes with Co^2+^ the Fe^2+^ ion necessary for the catalytic activity of PHD, hence inactivating the enzyme (Triantafyllou *et al.,* 2006; Elks *et al*., 2015; Munos-Sanchez and Chanez-Cardenas, 2018). As expected, wild type ESCs incubated for 24 hours with CoCl_2_ were characterized by high levels of expression of HIF1α target genes *Vegfa* and *Hk2* compared to untreated *Stat3^+/+^* cells. In contrast, *Stat3^-/-^* cells incubated with CoCl_2_ did not show significant upregulation of these transcripts (Fig. S2D,E), while *Hif1α* mRNA levels were not affected by the treatment. On the other hand, *Egln3*, which is a pure target gene of HIF1α involved in the elastic feedback loop of the oxygen deprivation response (Minamishima *et al.,* 2009), was not affected by the absence of *Stat3* (Fig. S2F,G). Interestingly, PLA analysis performed on ESCs treated with CoCl_2_ confirms the interaction between STAT3 and HIF1α (Fig. S2H). These results suggest that the formation of a nuclear STAT3-HIF1α complex is associated with full activation of hypoxia responsive genes.

### Hif1α transcriptional activity is not induced *in vivo* when Stat3 is knocked-out or inhibited

Zebrafish is an *in vivo* model in which the pathophysiological roles of HIF1α and JAK/STAT signalling have been extensively studied (Santhakumar *et al*., 2012; Gerri *et al.,* 2017; Gerri *et al*., 2018; Vettori *et al*., 2017; Marchi *et al*., 2020; Liu *et al.,* 2017; Peron *et al*., 2020; Peron *et al*., 2021; Dinarello *et al*., 2022). Thus, it appeared as a valid platform to analyze the STAT3-HIF1α crosstalk. We used the *Tg(4xHRE-TATA:mCherry,cmlc2:EGFP)^ia22^* hypoxia reporter zebrafish line (herein called HRE:mCherry), in which the mCherry red fluorescent protein is expressed in all tissues experiencing low oxygen tensions or pseudohypoxic conditions (Vettori *et al*., 2017). To investigate the requirement of Stat3 in the transcriptional response to hypoxia, we combined hypoxic and pseudohypoxic treatments with chemical or genetic inhibition of Stat3. To inhibit Stat3 signaling we used AG490, which blocks the Jak-mediated Y705 phosphorylation of Stat3 and abrogates its nuclear transcriptional activity (Park *et al.,* 2014; Peron *et al.,* 2020; Peron *et al*., 2021). Larvae treated with this compound were also characterized by a reduction of *stat3* gene expression (Fig. S3A). To induce hypoxia in zebrafish larvae, we incubated the animals for 3 days with 5% oxygen tension, while pseudohypoxia was forced by using either dimethyloxalylglycine (DMOG), an inhibitor of Phd-dependent degradation of Hif1α (Mole *et al*., 2003); CoCl_2_ or dexamethasone (Dex),a synthetic glucocorticoid that contributes to the stabilization of Hif1α by degrading Vhl (Vettori *et al*., 2017). Notably, while hypoxia and pseudohypoxia induced the fluorescence of HRE:mCherry zebrafish larvae, such induction was abolished by AG490, suggesting that, also in zebrafish, activated Stat3 is necessary for the correct induction of hypoxia-dependent transcription (Fig. 3A). Moreover, we tested the expression of *vegfa* and *hk1* at different times of exposure to low oxygen tension. We could observe that hypoxia determines a boost of expression of both *vegfa* and *hk1* after 8 hours of incubation in hypoxia; after 24 hours of incubation in hypoxic conditions, both transcripts return to the levels detected in normoxic larvae. Interestingly, *vegfa* expression increased after 72 hours of hypoxia, while *hk1* expression was upregulated after 48 hours of hypoxia (Fig. S3B,C). AG490 abrogated this hypoxia-dependent dynamical expression of *vegfa* and *hk1*, confirming the results obtained with the HRE:mCherry reporter line and with ESCs (Fig. S3B,C). STAT3 transcriptional activity has also been reported to be positively affected by the phosphorylation of Serine 727 (Levy *et al*., 2002; Zhang *et al*., 1995; Wen *et al*., 1995) and this post-translational modification is triggered by MEK/ERK pathway (Gough *et al*., 2013). As we recently demonstrated that the MEK/ERK inhibitor PD98059 blocks the pS727 activities of STAT3 (Peron *et al*., 2021), we sought to assess whether the inhibition of S727 phosphorylation can affect the responsiveness of larvae to low oxygen tensions. When HRE:mCherry reporter larvae are treated with PD98059 and incubated in low oxygen tensions, we could not observe significant differences in reporter fluorescence compared to hypoxic DMSO-treated larvae, suggesting that the chemical inhibition of S727 phosphorylation does not affect the transcriptional responsiveness of zebrafish to hypoxia (Fig. 3B). To further elucidate the role of Serine or Tyrosine phosphorylation in Stat3-dependent regulation of hypoxia transcription, we injected *Stat3* mRNAs (that we recently used and validated in Peron *et al*., 2021) in double transgenic eggs obtained from the breeding between *Tg(7xStat3-Hsv.Ul23:EGFP)^ia28^* and *HRE:mCherry* transgenic animals to monitor at the same time the Stat3- and the hypoxia-dependent transcription. Of note, wild type *Stat3* mRNA injection upregulated mCherry fluorescence suggesting that the overexpression of Stat3 increases hypoxia-dependent transcription, while *Stat3 Y705F* mRNA (that encodes a STAT3 that cannot be phosphorylated in 705 position, as described in Minami *et al*., 1996) cannot induce the fluorescence of both reporters (Fig. S3D,E). On the other hand, surprisingly, the overexpression of *Stat3 S727A* (in which the substitution of the Serine with an Alanine blocks the 727 phosphorylation, as shown in Wen *et al*., 1997), induces the Stat3-dependent transcriptional activity and the hypoxia-related gene expression (Fig. S3D,E). These results suggest that Tyrosine phosphorylation is necessary for the regulation of hypoxia by Stat3, while Serine is not.

**Fig. 3:**
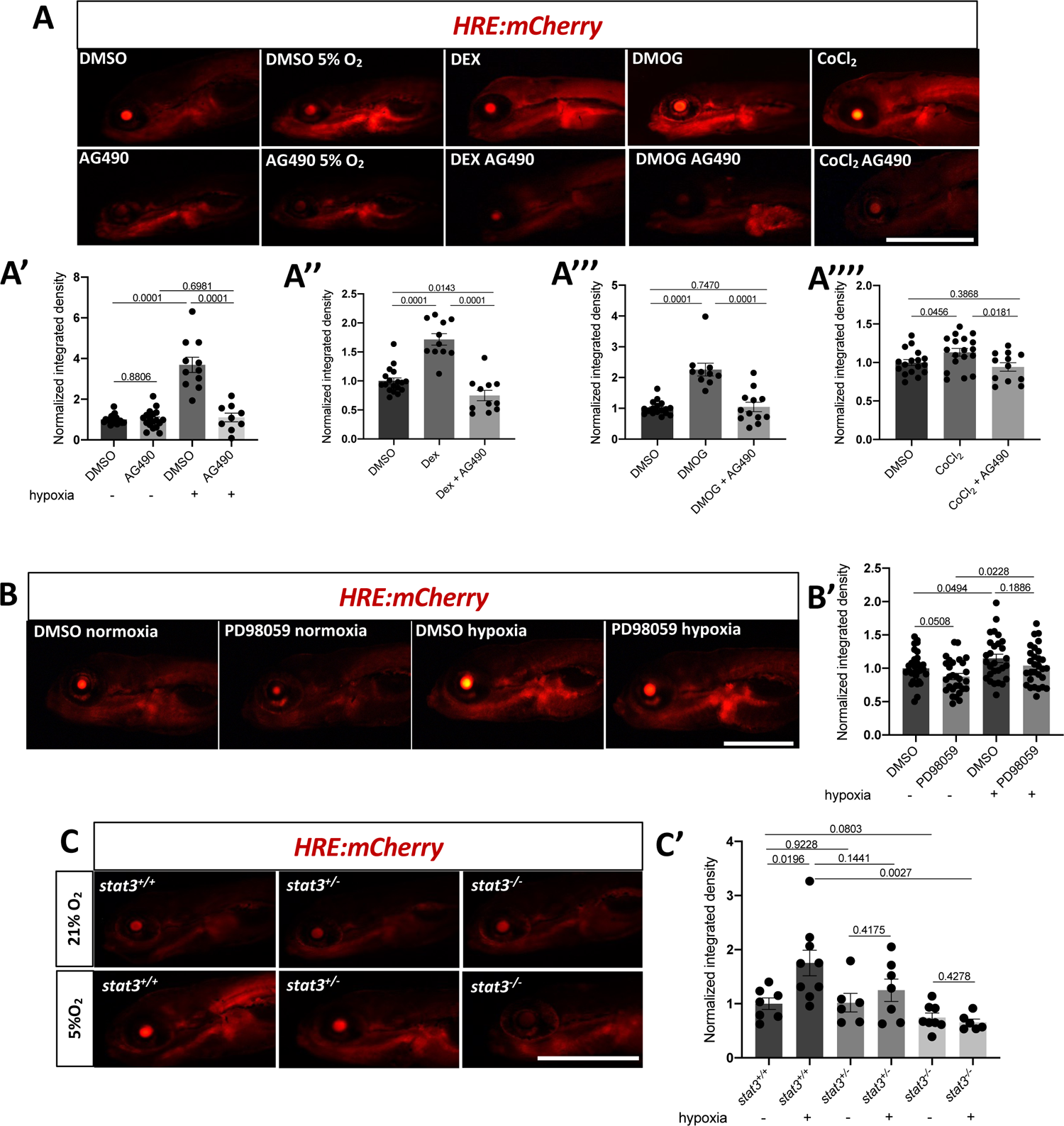
Stat3 is necessary for Hif1α transcriptional activity. A: Representative pictures of HRE:mCherry reporter zebrafish treated with AG490 in combination with low oxygen tension, DMOG, and Dex from 3-6 dpf and CoCl2 from 5-6 dpf. Scale bar 500 μm. A’: Fluorescence quantification of HRE:mCherry reporter zebrafish treated with AG490 in combination with low oxygen tension (n=10). A’’: Fluorescence quantification of HRE:mCherry reporter zebrafish treated with AG490 in combination with 10 μM Dex. A’’’: Fluorescence quantification of HRE:mCherry reporter zebrafish treated with AG490 in combination with 50 μM DMOG from 3-6 dpf. A’’’’: Fluorescence quantification of HRE:mCherry reporter zebrafish treated with AG490 in combination with 0.1 mM CoCl2 from 5-6 dpf. B: Representative pictures (B) and fluorescence quantification (B’) of HRE-mCHerry reporter zebrafish larvae treated with 12.5 mM PD98059 in combination with low oxygen tension. C: Representative pictures (C) and fluorescence quantification (C’) of *stat3^+/+^*, *stat3^+/-^* and *stat3^-/-^* sibling larvae in HRE:mCherry transgenic background treated with low oxygen tension from 3-6 dpf. Scale bar = 500 μm. Mean ± SEM. *p<0.05, **p<0.01, ***p<0.001, ****p<0.0001, ns=not significant.

Next, we crossed the *stat3^ia23^* mutant zebrafish line (herein called *stat3^-/-^*) (Peron *et al*., 2020) with the HRE:mCherry reporter line. We treated the HRE:mCherry*;stat3^+/+^*, HRE:mCherry*;stat3^+/-^* and HRE:mCherry*;stat3^-/-^* sibling larvae with 5% oxygen tension and measured mCherry fluorescence. Consistent with previous results obtained in ESC cells and in embryos treated with AG490, the Hif1α-dependent reporter activity increased in hypoxic *stat3^+/+^* larvae, while no significant increase of the reporter fluorescence was detected in hypoxic *stat3^+/-^ and stat3^-/-^* larvae when compared with normoxic siblings (Fig. 3C). These results indicate that the transcriptional activation of HIF1α targets requires STAT3 also *in vivo*.

### Nuclear crosstalk between Stat3 and Hif1α determines the regulation of hypoxia-dependent genes in zebrafish

Given that Stat3 is needed for the induction of normal Hif1α transcriptional activities and that hypoxia-induced mechanisms are impaired when Stat3 is inhibited or deleted, we wanted to determine by which mechanisms Stat3 regulates Hif1α transcriptional activity.

We first tested whether Stat3 inactivation would affect *hif1α* expression levels, its stabilization, or the levels of HIF1α regulators *vhl* and *egln3;* we observed no significant differences in their expression levels when comparing *stat3^+/+^* and *stat3^-/-^* 6-dpf larvae (Fig. S4A-D). We conclude that, as already observed in ESCs (Fig. 2B), the lack of a functional Stat3 does not impair the expression and stabilization of Hif1α in zebrafish larvae.

Given that in ESCs, STAT3 and HIF1α interact in the nucleus, we focussed our attention on the nuclear activity of HIF1α *in vivo*. For this purpose, we injected an mRNA that encodes a dominant active (DA) form of *hif1αb* bearing mutations at two prolines and one asparagine (P402A, P564G, N804A), hence preventing their hydroxylation by Phd and subsequent Vhl-dependent degradation (Elks *et al*., 2011; Elks *et al*., 2013). In other words, this construct activates by default nuclear targets, independently from any upstream degrading cue. We treated HRE:mCherry hypoxia reporter larvae injected with *hif1αb DA* mRNA at 3 dpf with AG490 and analysed the reporter fluorescence at 4 dpf. Notably, *hif1αb DA* mRNA determines an induction of the reporter activity that is significantly decreased in injected larvae treated with AG490 (Fig. 4A). We further investigated the endogenous genes induced by *hif1αb DA* mRNA by performing RT-qPCR analysis. The expression levels of Hif1α-dependent genes, such as *vegfa* and *hk1* were induced by *hif1αb DA* mRNA, however, AG490 abrogated their upregulation (Fig. 4B,C). These results confirmed *in vivo* that activation of HIF1α nuclear targets rely on active STAT3, as observed in mouse ESCs.

**Fig. 4:**
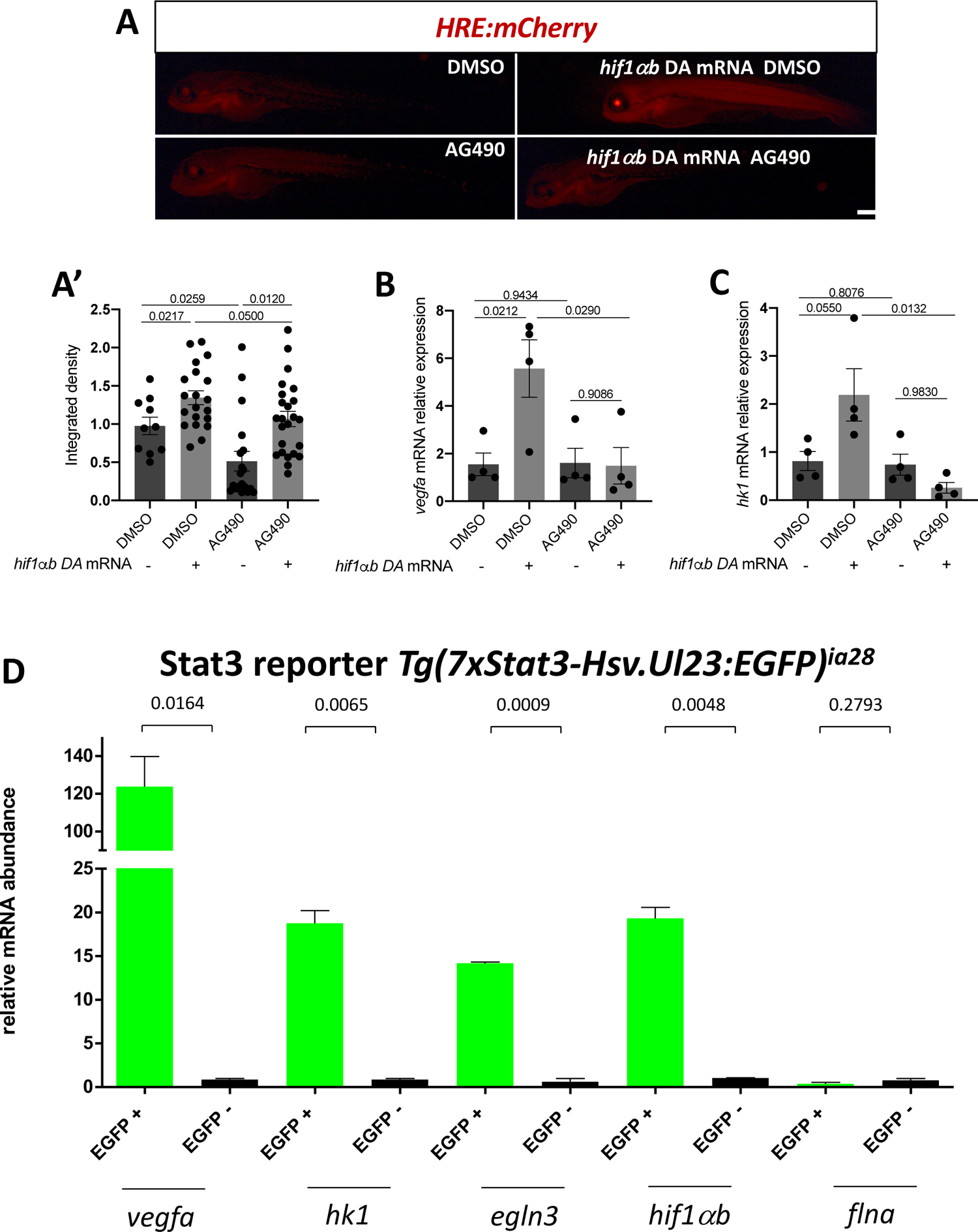
Stat3 and Hif1α cooperate in the nucleus for the transcription of Hif1α target genes. A: representative pictures (A) and fluorescence quantification (A’) of 4-dpf HRE:mCherry reporter larvae treated with either DMSO or 50 μM AG490 for 24 hours after injection of *hif1ab* DA mRNA (n = 3 independent biological replica, each dot represent a larva). B: RT-qPCR analysis of *vegfa* of 4-dpf larvae injected with *hif1ab* DA mRNA and treated with AG490 (n = 4 independent biological replicates composed by pool of 15 larvae, each dot represents a pool of larvae). C: RT-qPCR analysis of *hk1* of 4-dpf larvae injected with *hif1αb* DA mRNA and treated with AG490 (n = 4 independent biological replicates composed by pool of 15 larvae, each dot represents a pool of larvae). D: RT-qPCR analysis of *vegfa*, *hk1*, *egln3*, and *hif1αb* on EGFP-positive and EGFP-negative cells sorted from *ia28* transgenic adult intestines (n=3). Mean±SEM. *p<0.05; **p<0.01; ***p<0.001; ns = not significant.

Additionally, we took advantage of the zebrafish model to ask whether HIF1α and STAT3 are active in the same cells (cell autonomously), or whether they co-operate non-cell-autonomously in distinct cells somehow interconnected. To do so, we used the Stat3 reporter line *Tg(7xStat3-Hsv.Ul23:EGFP)^ia28^* (Peron *et al*. 2020), and sorted Stat3 responsive intestinal stem cells. Results of RT-qPCR revealed a strong and significant upregulation of Hif1α transcriptional activity in EGFP sorted cells. In particular, *hif1αb* transcript levels were 20-fold higher in EGFP-positive cells compared to EGFP-negative cells. The upregulation of the *vegfa* and *hk1,* involved in angiogenesis and glucose metabolism, as well as of the pure Hif1α target gene *egln3*, involved in the Hif1α degradation feedback loop (Minamishima *et al*., 2009), revealed a massive Hif1α transcriptional activity in the EGFP-positive Stat3-responsive cells (Fig 4D). *flna*, a gene that is neither affected by Stat3 nor by Hif1α was used in this experiment as a negative control. We conclude that in the context of a complex tissue like the intestine, HIF1α direct targets are strongly expressed in cells with active STAT3, further indicating cooperative transcriptional activation by the two factors and the cell-autonomous regulation of the STAT3-HIF1α crosstalk.

### Hypoxia-dependent processes are impaired in *stat3* mutant zebrafish

Our results so far indicate a cooperative activation of STAT3 and HIF1α in the transcriptional response to hypoxia. We then asked whether such cooperation is important also for the physiological responses in an organism.

One of the most relevant processes induced by hypoxia is angiogenesis (Fraisl *et al*., 2009). To study how this process is affected by hypoxia in *stat3* mutant zebrafish larvae, we decided to use the zebrafish endothelial cell reporter line *Tg(Fli1:EGFP)^y1^* (Lawson and Weinstein, 2002). *stat3^+/+^*, *stat3^+/-^* and *stat3^-/-^* sibling larvae in *Tg(Fli1:EGFP)^y1^* transgenic background were treated with 5% oxygen tension from 48 hpf to 54 hpf. After the treatment, we detected a significant increase of endothelial fluorescence in *stat3^+/+^* hypoxic larvae when compared with untreated siblings (Fig. 5 A), indicating that angiogenesis is induced by low oxygen tensions, hence confirming what already observed by Eyries *et al*. (2008). A similar induction of fluorescence was also detected in hypoxic heterozygous *stat3^+/-^* larvae when compared to normoxic heterozygous siblings, but no significant differences were detected between normoxic and hypoxic null *stat3^-/-^* larvae, indicating a pivotal role of Stat3 in this process (Fig. 5 A).

**Fig. 5:**
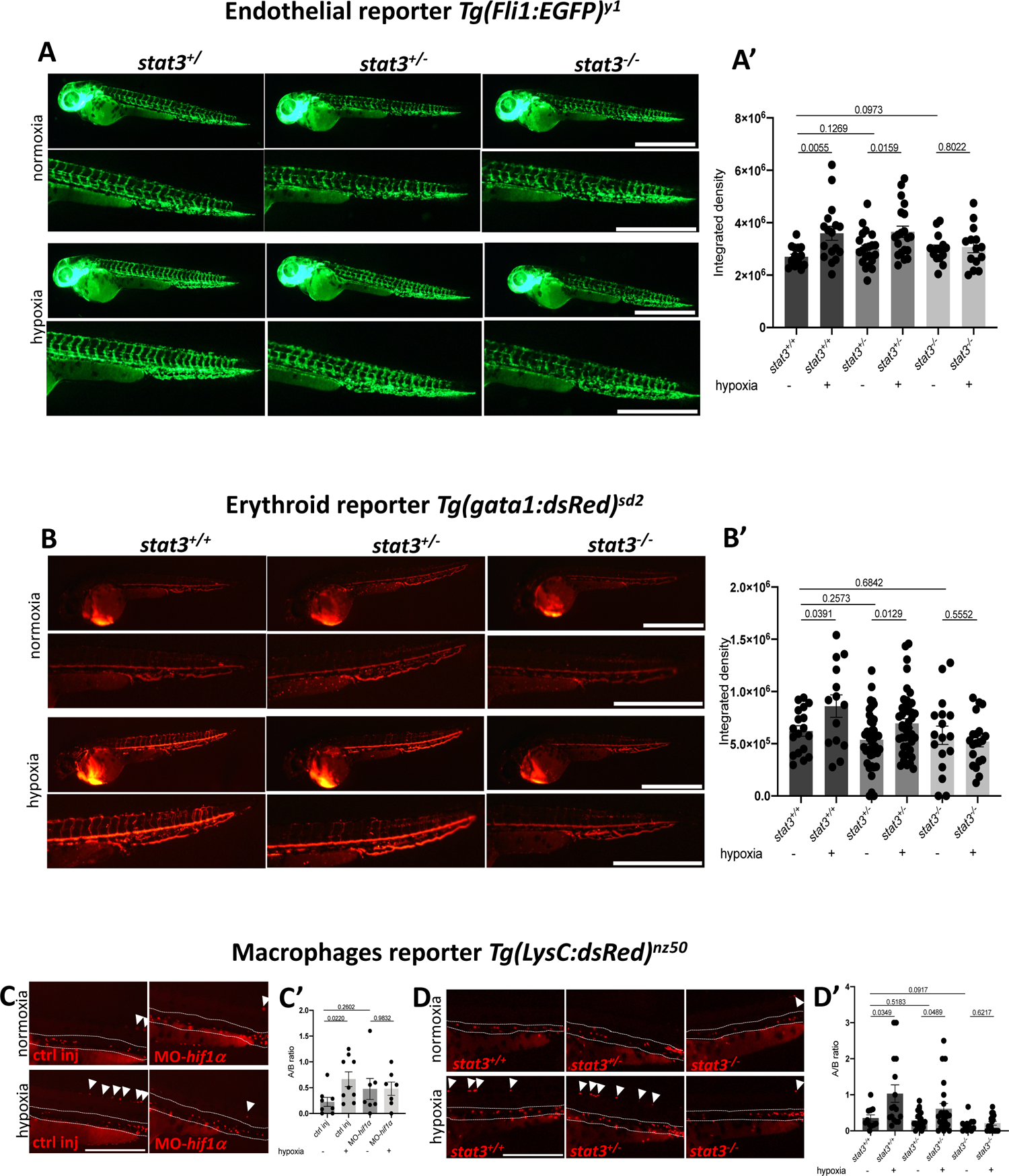
*stat3* genetic ablation affects angiogenesis, and macrophage migration. A: Representative pictures (A) and fluorescence quantification of the trunk (A’) of 54-hpf *stat3^+/+^*, *stat3^+/-^* and *stat3^-/-^* in *Tg(Fli1:EGFP)^y1^* transgenic background incubated in normoxia and hypoxia for 6 hours. Scale bar: 1 mm. B: Representative pictures. (B) and fluorescence quantification of the tail (B’) of 54-hpf *stat3^+/+^*, *stat3^+/-^* and *stat3^-/-^* larvae in *Tg(gata1:dsRed)^sd2^* transgenic background incubated in normoxia and hypoxia. Scale bar: 1 mm. C: Representative pictures of 54-hpf control (ctrl inj) and *hif1α* morphants (MO-*hif1α*) in *Tg(LysC:dsRed)^nz50^* transgenic background incubated in normoxia and hypoxia for 6 hours (scale bar: 500 μm); quantification of the ratio of dsRed-positive cells in region A and in region B (C’). Area B between dashed lines. Arrowheads point at fluorescent cells in Area A. D: Representative pictures of 54-hpf *stat3^+/+^*, *stat3^+/-^* and *stat3^-/-^* larvae in *Tg(LysC:dsRed)^nz50^* transgenic background incubated in normoxia and hypoxia for 6 hours (scale bar: 500 μm); quantification of the ratio of dsRed-positive cells in region A and in region B (D’). Area B between dashed lines. Arrowheads point at fluorescent cells in Area A. Mean±SEM. *p<0.05; **p<0.01; ***p<0.001; ns=not significant.

Hypoxia plays a central role in erythropoiesis (Haase, 2010; Zhang *et al.,* 2012; Solak *et al*., 2016; Kietzmann, 2020) and in zebrafish *gata1* is a specific marker for erythropoiesis and erythrocytes (Lyons *et al*., 2002; Galloway *et al*., 2005; Bresciani *et al*., 2010; Quintana *et al*., 2014; Li *et al*., 2014; Lenard *et al*., 2016). Hence, we decided to use the *Tg(gata1:dsRed)^sd2^* transgenic line (Galloway *et al*., 2005), in which erythroid cells display a strong red fluorescence, to see whether Stat3 has a role in hypoxia-induced erythropoiesis.

Notably, 6-hour long hypoxia treatments determine a significant increase of fluorescence in *stat3^+/+^*. Similarly, an upregulation of fluorescence was detected also in hypoxic *stat3^+/-^* larvae. Of note, no significant differences were detected between normoxic and hypoxic *stat3^-/-^* larvae, demonstrating that Stat3 is involved in the hypoxia-induced erythropoiesis (Fig. 5B).

Since the hypoxia/HIF1 pathway has been linked to mobilization and polarization of macrophages (Lewis *et al.,* 1999; Elks *et al*., 2013; Ke *et al*., 2019; Lewis and Elks, 2019; Sadiku and Walmsley, 2019), we used the *Tg(LysC:dsRed)^nz50^* transgenic line (Hall *et al*., 2007), to focus our attention on them. First, to confirm that the migration of macrophages outside the aorta-gonad-mesonephros (AGM) region is triggered by hypoxia and relies on Hif1α, we injected 1-cell stage *Tg(LysC:dsRed)^nz50^* transgenic embryos with a solution containing both morpholinos (MO) against *hif1αa* and *hif1αb* (as previously described in Gerri *et al*., 2017). 48-hpf controls and *hif1α* morphants were subsequently incubated in normoxia or hypoxia for 6 hours and the number of macrophages was counted. As reported in Fig. 5C and Fig. S5A, hypoxia determines an increase of the ratio between the number of cells observed in the trunk of larvae (named region A) and that in the AGM (named region B), suggesting that hypoxia determines the migration of macrophages away from AGM. Interestingly, we could not detect significant differences between normoxic and hypoxic morphants, demonstrating that Hif1α is required for this process. Subsequently, we sought to see whether Stat3 is also involved in this process. To do so, we treated 48-hpf *stat3^+/+^*, *stat3^+/-^* and *stat3^-/-^* larvae *Tg(LysC:dsRed)^nz50^* transgenic background with 5% oxygen tension for 6 hours. Treated *stat3^+/+^* and *stat3^+/-^* showed a significant increase of Area A/Area B ratio compared to untreated siblings, while hypoxia did not have any effect in mobilizing macrophages of *stat3^-/-^* larvae, suggesting that Stat3 is involved in this Hif1α-dependent process (Fig. 5D and Fig. S5B). Moreover, we analysed the level of expression of genes involved in macrophage activity, like *mfap4*, *tek,* and *lcp1* (Walton *et al*., 2020; Melcher *et al.,* 2008; Kell *et al*., 2018). *mfap4* and *tek* appeared to be downregulated in their expression in *stat3^-/-^* when compared to *stat3^+/+^* siblings (Fig. S5C), highlighting a role of Stat3 in the homeostasis of these cells. Importantly, the total number of dsRed-positive cells does not seem to be affected significantly by genetic ablation of *stat3,* indicating a specific effect on migration rather than survival or proliferation (Fig. S5B,C).

## DISCUSSION

In the last 20 years, several research groups suggested an interplay between STAT3 and HIF1: for instance, Gray and collaborators in 2005 demonstrated that the expression of *Vegfa* in pancreatic and prostate carcinomas depends on HIF1α, STAT3, CBP/p300 and Ref-1/APE (Gray *et al*., 2005) and this result was further investigated by Oh and collaborators (2011), who showed that STAT3 and HIF1α bind the *Vegfa* promoter (Oh *et al*., 2011). However, up to now, no clear description of STAT3-HIF1α connections have been provided yet and the mechanisms through which STAT3 and HIF1α regulate each other is still controversial: some groups demonstrated that STAT3 triggers the expression of *HIF1α* mRNA (Jung *et al*., 2005; Cui *et al*., 2016), others suggested that STAT3 is involved in the stabilization of HIF1α protein (Xu *et al.,* 2005). To better elucidate this intricate mechanism of regulation, we decided to analyze the crosstalk between STAT3 and HIF1α using both *in vivo* and *in vitro* physiological models. RNAseq experiments performed on mouse ESCs allowed us to determine that a large part (about 30%) of genes induced by hypoxia and Hif1α require a functional STAT3. Of note, with this experiment we provide the list of genes that are regulated in concert by STAT3 and HIF1α, demonstrating that the hypoxia-triggered control of glycolysis and angiogenesis depends on STAT3.

Taking advantage of the Stat3 zebrafish reporter line characterized in Peron *et al*. (2020), we demonstrated that Stat3-positive cells in the zebrafish intestine have a strong Hif1α signature, hence, we decided to test whether hypoxia-dependent activities rely on Stat3. Drugs inhibiting Stat3 signaling pathway established a severe unresponsiveness of zebrafish larvae to hypoxia and pseudohypoxia, underlying a pivotal role for Stat3 in the normal induction of hypoxia-related transcriptional activity. This result was additionally confirmed by the defective responsiveness to hypoxia of *stat3* mutant zebrafish larvae: many hypoxia- and Hif1α-dependent processes like angiogenesis and macrophage mobilization from AGM are significantly affected by *stat3* genetic ablation, confirming our hypothesis that Stat3 is involved in the correct activation of hypoxic adaptation. Notably, we could show both in zebrafish and mouse ESCs that STAT3 is not involved in the transcription of *Hif1α* gene, nor in the stabilization of HIF1α protein and that HIF1α localizes in the nucleus even when *Stat3* is not expressed. Remarkably, we demonstrated that STAT3 physically interacts with HIF1α and that their interaction is specifically localized in the nucleus. These results support the hypotheses that STAT3 enhances HIF1α transcriptional activities by regulating its interaction with either HIF1β, the Hypoxia Responsive Elements on target genes or the basal transcriptional machinery. Finally, it is worth mentioning that STAT3-dependent control of HIF1α transcriptional activity affects specifically genes involved in angiogenesis (*Vegfa*) and metabolism (*Hk1, Hk2, Pfkp*) and this result was observed both in mouse ESCs (Fig 1) and in zebrafish (Fig. 4, 5). The regulation of these processes, representing two of the most important biological functions regulated by HIF1α, highlights the pivotal role of STAT3 in the proper response to low oxygen tensions and shows that the connection between STAT3 and HIF1α is much stronger than it was previously thought. These findings, together with the observation by Russell *et al*. (2011) that VHL promotes ubiquitin-mediated destruction of JAK2, and the fact that JAK/STAT3 and HIF/VHL play a role in many diseases, call for a deeper analysis of their crosstalk and the implications this crosstalk might have from a therapeutic point of view.

## MATERIAL AND METHODS

### Animals husbandry and zebrafish lines

Animals were staged and fed as described by Kimmel *et al*. (1995) and maintained in a large-scale aquaria system.

Embryos were obtained after natural mating, raised in Petri dishes containing fish water (50x: 5 g NaHCO_3_, 39.25 g CaSO_4_, 25 g Instant Ocean for 1 l) and kept in a 12:12 light dark cycle at 28° C. All experimental procedures complied with European Legislation for the Protection of Animals used for Scientific Purposes (Directive 2010/63/EU). *stat3^ia23^* mutant line (Peron *et al*., 2020) is genotyped by PCR amplification and 3% agarose gel migration. The *Tg(7xStat3-Hsv.Ul23:EGFP)^ia28^* line, the *Tg(4xHRE-TATA:mCherry,cmlc2:EGFP)^ia22^* line, the *Tg(fli1:EGFP)^y1^,* the *Tg(gata1:dsRed)^sd2^* line and the *Tg(LysC:dsRed)^nz50^* lines have been respectively characterized by Peron *et al*. (2020), Vettori *et al*. (2017), by Lawson and Weinstein (2002), Galloway *et al*. (2005) and by Hall *et al*. (2007). All animal experiments were performed under the permission of the ethical committee of the University of Padova and the Italian Ministero della Salute (23/2015-PR).

### Mouse ESCs culture and treatments

ESCs were grown as described in Betto *et al*. (2021). In detail, wild type (*Stat3^+/+^*) or *Stat3* knock-out (*Stat3^-/-^*) (described previously in Ying *et al*. (2008), Carbognin *et al*. (2016), Takeda *et al*. (1997) and provided by A. Smith’s laboratory) mouse ESC lines were routinely cultured without feeders on gelatin-coated plates (0.2% gelatin, Sigma-Aldrich, cat. G1890). Media were changed every 2 days and cells were passaged when approaching confluency (every 2-3 days); to passage, cells were replated at required density following dissociation with Accutase (ThermoFisher, cat. A1110501). Cells were grown in 2iLIF culture conditions, prepared as follows: serum-free KSR (Knockout Serum Replacement) 10% (ThermoFisher, cat. 10828028) - based medium in GMEM (Sigma-Aldrich, cat. G5154) supplemented with 1% FBS (ThermoFisher, cat. 10270106), 100 mM 2-mercaptoethanol (Sigma-Aldrich, cat. M3148), 1× MEM non-essential amino acids (ThermoFisher, cat. 11140050), 2 mM L-glutamine (ThermoFisher, cat. 25030081), 1 mM sodium pyruvate (ThermoFisher, cat. 11360070), and with small-molecule inhibitors PD (1 μM, PD0325901), CH (3 μM, CHIR99021) from Axon (cat. 1386 and 1408) and LIF (100 units/ml, Qkine, cat. Qk036).

All cells were maintained at 37°C in humidified incubators with 5% CO_2_. Hypoxia treatments were performed for 24 hours with 1% oxygen tensions. 0.1 mM CoCl_2_ was added to cell culture media for 24 hours.

### Proximity ligation assay

Mouse ESCs were fixed for 10 minutes in 4% formaldehyde at −20°C, washed in TBS and permeabilized for 10 minutes with TBST + 0.5% Triton X-100 at room temperature. PLA was performed following the Duolink® protocol: cells were covered with Duolink® blocking solution and incubated for 60 minutes at 37°C in humidity chamber; primary antibodies anti-STAT3 mouse monoclonal (Cell Signaling, 9139) (1:100) and anti-HIF1α rabbit monoclonal (Novus Biologicals, NB100-499) (1:100) were dissolved in Duolink® antibody diluent and samples were incubated overnight at 4°C. Primary antibodies solution was removed and samples were washed 2 times for 5 minutes with washing buffer A (prepared following manufacturer’s instructions) and subsequently incubated for 1 hour at 37°C with Duolink® PLA probe mix. After 2 washes with washing buffer A for 5 minutes each, samples were incubated at 30 minutes at 37°C with Duolink® ligation mix. Samples were washed 2 times for 2 minutes with washing buffer A and, subsequently, incubated for 100 minutes at 37°C with Duolkin® amplification mix. After three washes with washing buffer B (prepared following manufacturer’s instructions), samples were ready for imaging.

### Drug and hypoxic treatments

We used the following chemical compounds: AG490 (T3434, Sigma); DMOG (D3695, Sigma), Dex (D1756, Sigma), CoCl_2_ (232696, Sigma). Dex was diluted in ethanol and stored at 4° C, while the other compounds were diluted in DMSO and stored in small aliquots at −20° C. 2 mM 1-phenyl-2-thiourea (PTU) was used to inhibit pigmentation. Larvae were treated from 3 dpf to 6 dpf with 50 μM AG490, 50 μM DMOG, 10 μM Dex; from 5 dpf to 6 dpf with 0.5 mM CoCl_2_. 5% oxygen tension was maintained using ProCO_2_ (BioSpherix) device and larvae were incubated with low oxygen tension from 3 dpf to 6 dpf.

### mRNA isolation and quantitative real time reverse transcription PCR (qRT-PCR)

Total RNAs were extracted from pools of 20 larvae at 3, 4, 5 and 6 dpf with TRIzol reagent (Thermo Fisher Scientific, 15596018) and incubated at 37° C for 30 minutes with RQ1 RNase-Free DNase (Promega, M6101). cDNA synthesis was performed using random primers (Promega, C1181) and M-MLV Reverse Transcriptase RNase H (Solis BioDyne, 06-21-010000) according to the manufacturer’s protocol. qPCRs were performed in triplicate with CybrGreen method by means of Rotor-gene Q (Qiagen) and the 5x HOT FIREPol EvaGreen qPCR Mix Plus (Solis BioDyne, 08-36-00001) and *zgapdh* and *mActnb* were used as internal standard in each sample. The amplification protocol consists of 95° C for 14 minutes followed by 45 cycles at 95° C for 20 seconds, 60° C for 20 seconds and 72° C for 25 seconds. Threshold cycles (Ct) and melting curves were generated automatically by Rotor-Gene Q series software and results were obtained with the method described in Livak and Schmittgen (2001). Sequences of genes of interest primers were designed with Primer3 software (Untergasser *et al*., 2012) (http://bioinfo.ut.ee/primer3-0.4.0/input.htm) and are listed in supplementary material Table 1.

**Table 1:**
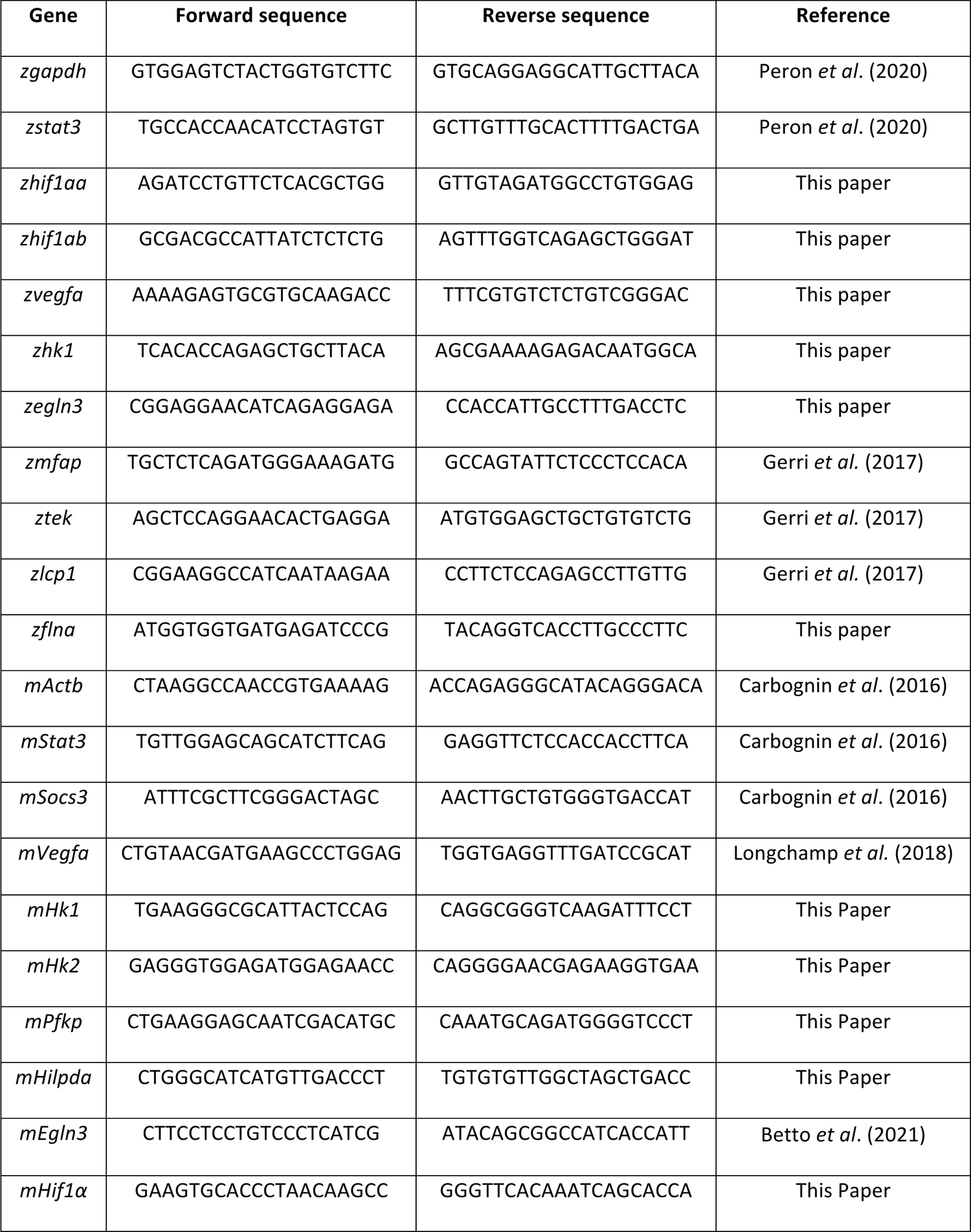
list of primers used for RT-qPCR (5’ → 3’ sequences).

### Protein extraction and western blotting

Total protein extracts were obtained by homogenization of pools of 20 6-dpf larvae in ice cold RIPA buffer (ThermoFisher, 89900) and Complete EDTA-free protease inhibitor cocktail (Sigma, 11873580001). For western blot analysis 40 μg of protein extracts were loaded per well on Bolt 4-12% Bis-Tris Plus Gels (ThermoFisher, NW04120BOX) and blotted on PVDF immobilon-p membranes (Millipore, IPFL00010). Dried membranes were then washed with PBS (Sigma, P4417) with 0.1% (w/v) Tween20 and incubated overnight with primary antibodies at 4° C: anti-HIF1α (1:500, MA1-16504 Invitrogen); anti-STAT3 (1:1000, 9139S Cell Singaling); anti-βActin (1:5000, MA1-744 ThermoFisher). Secondary anti-Rabbit and anti-Mouse HRP-conjugated antibodies (170-6515 BIORAD, and 170-6516 BIORAD, respectively) were incubated for 1 hour at room temperature and protein bands detected by chemiluminescence on an Alliance MINI HD 9 Blot Imaging System. Quantification of the signal was performed with ImageJ.

### Imaging

For *in vivo* imaging, transgenic larvae were anesthetized with 0.04% tricaine, embedded in 1% low-melting agarose and mounted on a depression slide. Nikon C2 confocal system was used to acquire images from *Tg(Fli1:EGFP)^y1^* transgenic larvae. *Tg(4xHRE-TATA:mCherry,cmlc2:EGFP)^ia22^, Tg(LysC:dsRed)^nz50^* and *Tg(gata1:dsRed)^sd2^* transgenic larvae were mounted in 1% low-melting agarose and observed with a Leica M165 FC microscope equipped with a Nikon DS-Fi2 digital camera. All images were analysed with Fiji (ImageJ) software and fluorescence integrated density was calculated setting a standard threshold on non-fluorescent samples.

### Fluorescence-Activated Cell Sorting (FACS)

Adult intestines of *Tg(7xStat3-Hsv.Ul23:EGFP)^ia28^* fish were dissected and treated as reported by Peron *et al*. (2020). Cells from two intestines were pooled together for three independent biological replicas.

### In vitro mRNA synthesis and mRNA microinjection

The DA *hif1αb* mRNA was obtained from pCS2-*hif1abDA* vector that Dr. Phil Elks kindly sent us. mRNAs were *in vitro* transcribed using the mMESSAGE mMACHINE® SP6 Transcription Kit (Thermo Fisher Scientific) and purified using the RNA Clean and Concentrator kit (Zymo Research). A mix containing 50ng/μM mRNA Danieu injection Buffer and Phenol Red injection dye, was injected into 1-cell stage embryos. *hif1αa* and *hif1αb* morpholinos, that Dr. Nana Fukuda kindly sent us, were injected together in 1-cell stage embryos as described in Gerri *et al*., 2017.

### RNA-sequencing

Quant Seq 3’ mRNA-seq Library Prep kit (Lexogen) is used for library construction. Library generation is initiated by oligodT priming. The primer already contains Illumina-compatible linker sequences. After first strand synthesis the RNA is removed and second strand synthesis is initiated by random priming and a DNA polymerase. The random primer also contains Illumina-compatible linker sequences. Second strand synthesis is followed by a magnetic bead-based purification step. The library is then amplified, introducing the sequences required for cluster generation. External barcodes are introduced during the PCR amplification step. Library quantification is performed by fluorometer (Qubit) and bioanalyzer (Agilent). QuantSeq Forward contains the Read 1 linker sequence in the second strand synthesis primer, hence NGS reads are generated towards the poly(A) tail and directly correspond to the mRNA sequence. QuantSeq FWD maintains strand-specificity and allows mapping of reads to their corresponding strand on the genome, enabling the discovery and quantification of antisense transcripts and overlapping genes. Sequencing is performed on NextSeq500 ILLUMINA instrument to produce 5 million of reads (75bp SE) for sample. The reads were trimmed using BBDuk (BBMap v. 37.62), with parameters indicated in the Lexogen data analysis protocol. After trimming, reads were aligned to the mouse genome (GRCm38.p6) using STAR (v. 2.7.6a). The gene expression levels were quantified using featureCounts (v. 2.0.1). Genes were sorted removing those that had a total number of counts below 10 in at least 4 samples out of 16. After applying this filter, we identified 12,432 expressed genes that were considered for further analyses. All RNA-seq analyses were carried out in R environment (v. 4.0.0) with Bioconductor (v. 3.7). We computed differential expression analysis using the DESeq2 R package (v. 1.28.1) (Love *et al*., 2014 Genome Biol). DESeq2 performs the estimation of size factors, the estimation of dispersion for each gene and fits a generalized linear model. Transcripts with absolute value of log2[FC] > 0.75 and an adjusted p-value < 0.01 (Benjamini–Hochberg adjustment) were considered significant and defined as differentially expressed for the comparison in the analysis. Heatmaps were made using counts-per-million (CPM) values with the pheatmap function from the pheatmap R package (v.1.0.12; distance = ‘correlation’, scale = ‘row’) on differentially expressed genes or selected markers. Volcano plots were computed with log2[FC] and −log10[q-value] from DESeq2 differential expression analysis output using the ggscatter function from the ggpubr R package (v. 0.4.0).

### Statistical analysis

Statistical analysis was performed using Graph Pad Prism software V6.0. Data are presented as the means ± SEM. Comparison between different groups of samples was performed by Student’s t-test with a confidence interval of 95%. The p-values are indicated with asterisks: *p< 0.05; **p< 0.01; ***p< 0.001; ****p<0.0001.

## ACKNOWLEDGMENTS

We are thankful to the personnel at the Zebrafish Centre of the University of Padova (Luigi Pivotti, Shkendy Iljazi, and Martina Milanetto). We are thankful to Valentina Tonelotto, Davide Volpato, Lorenzo Badenetti, Giada Vanni, Lorenzo Lupi, Saverio Fortunato and Olga Perrotti for their technical support. The study has been supported by the AIRC project IG-2017-19928 to F.A. G.M.’s laboratory is supported by grants from the Giovanni Armenise-Harvard Foundation, the Telethon Foundation and an ERC Starting Grant (MetEpiStem).

## Authors’ contributions

AD acquisition, analysis, interpretation of data, drafting the work, writing-original draft; RMB acquisition, analysis, interpretation of data, drafting the work, writing-review & editing; CC acquisition, analysis, interpretation of data, writing-review & editing; LD acquisition, analysis, interpretation of data; GMe acquisition, analysis, interpretation of data; MP acquisition, analysis; AT acquisition, analysis; RG acquisition, analysis; CL acquisition, analysis; NT funding management; GMa interpretation of data, drafting the work, writing-review & editing; FA interpretation of data, drafting the work, writing-review & editing.

## Supplementary figures

**Fig. S1:**
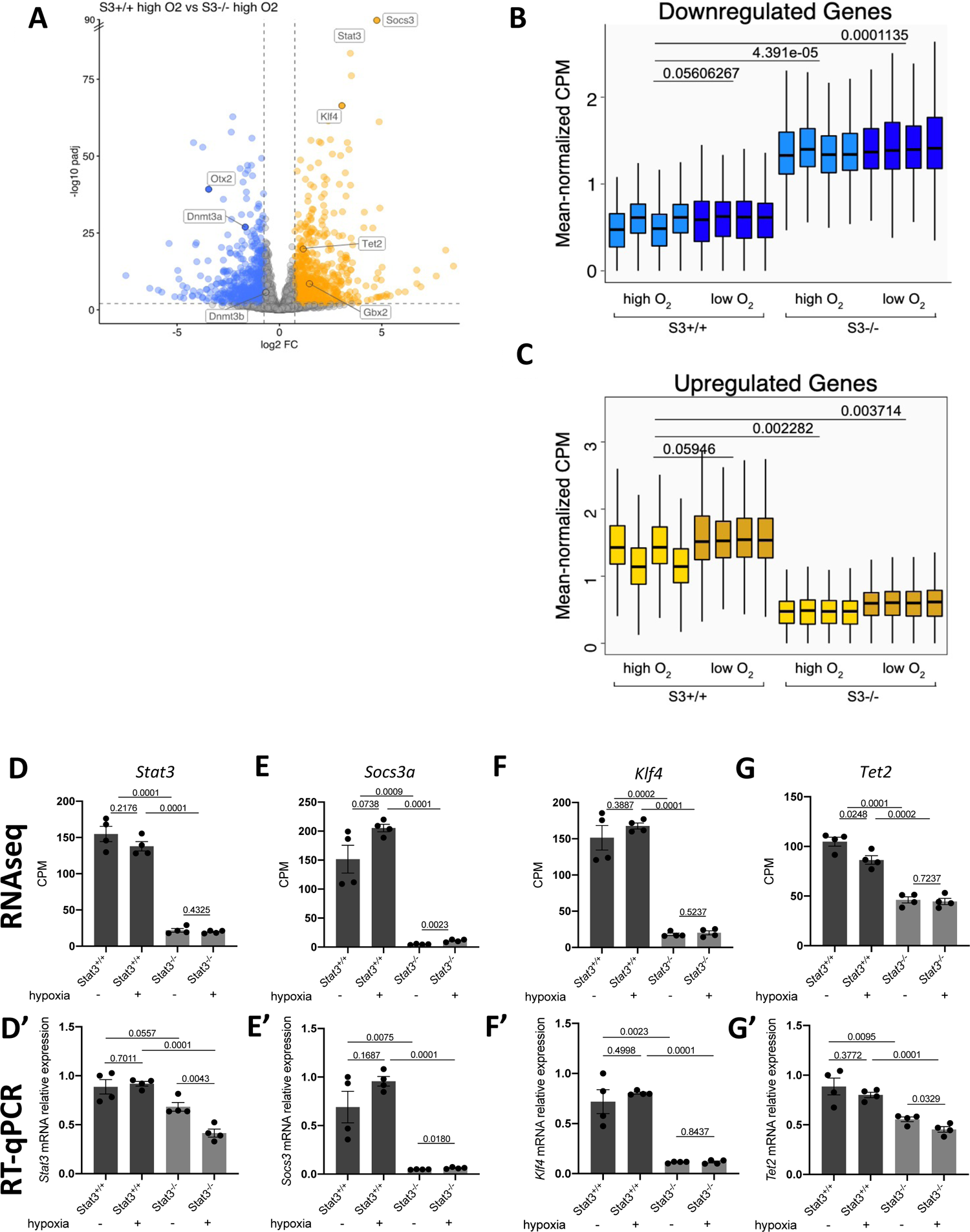
Transcriptome analysis by RNA-seq. A: Genes that were differentially expressed (log2[fold change (FC)] > + 0.75 or < −0.75; q-value < 0.01, Benjamini–Hochberg adjustment, as indicated by dashed lines) between *Stat3^+/+^* and Stat3*^-/-^* cells; n = 4 biological replicates. B: Expression levels of genes that were downregulated in *Stat3^+/+^* relative to *Stat3^-/-^* cells. C: Expression levels of genes that were upregulated in *Stat3^+/+^* relative to *Stat3^-/-^* cells. Each boxplot shows the 1st, 2nd and 3rd quartiles; the whiskers show the minimum and maximum values. D: *Stat3* expression in *Stat3^+/+^* and *Stat3^-/-^* cells incubated either in normoxia and hypoxia measured with RNAseq (D) and RT-qPCR (D’). E: *Socs3* expression in *Stat3^+/+^* and *Stat3^-/-^* cells incubated either in normoxia and hypoxia measured with RNAseq (E) and RT-qPCR (E’). F: *Klf4* expression in *Stat3^+/+^* and *Stat3^-/-^* cells incubated either in normoxia and hypoxia measured with RNAseq (F) and RT-qPCR (F’). G: *Tet2* expression in *Stat3^+/+^* and *Stat3^-/-^* cells incubated either in normoxia and hypoxia measured with RNAseq (G) and RT-qPCR (G’). Mean=SEM. **p<0.01, ***p<0.001, ns=not significant.

**Fig. S2:**
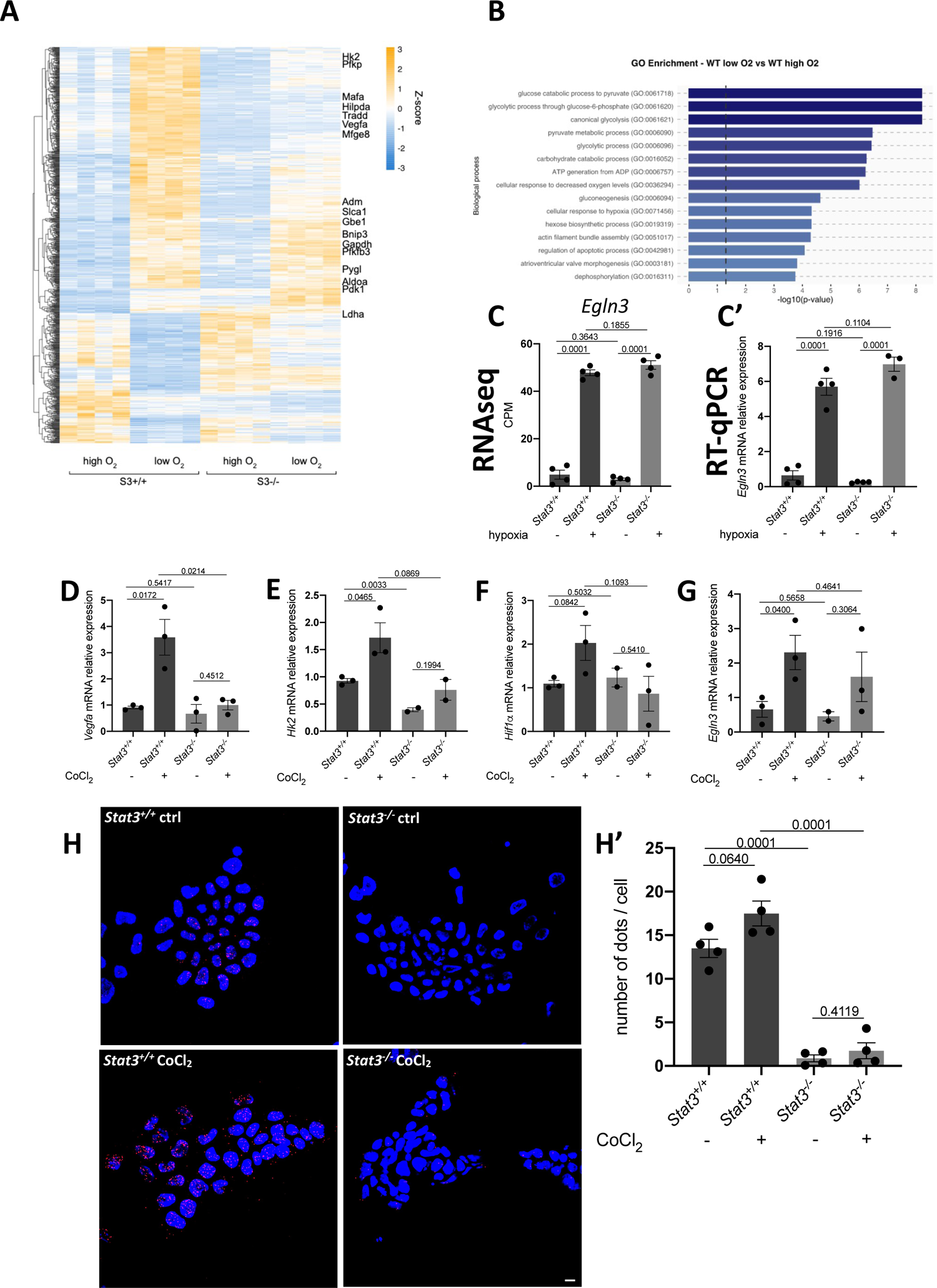
hypoxia and CoCl2 induce HIF1α target genes in *Stat3^+/+^* cells. A: heatmap of RNAseq data reporting the expression of genes that are affected by hypoxia in *Stat3^+/+^* cells compared to *Stat3^+/+^* normoxic cells. B: GO enrichment between normoxic and hypoxic *Stat3^+/+^* cells. C: expression level of *Egln3* in normoxic and hypoxic *Stat3^+/+^* and *Stat3^-/-^* mESCs obtained from RNAseq data (C) and from RT-qPCR (C’). D: RT-qPCR analysis of *Vegfa* mRNA expression in *Stat3^+/+^* and *Stat3^-/-^* ESCs incubated either with DMSO or CoCl2. E: RT-qPCR analysis of *Hk2* mRNA expression in *Stat3^+/+^* and *Stat3^-/-^* ESCs incubated either with DMSO or CoCl2. F: RT-qPCR analysis of *Hif1α* mRNA expression in *Stat3^+/+^* and *Stat3^-/-^* ESCs incubated either with DMSO or CoCl2. G: RT-qPCR analysis of *Phd3* mRNA expression in *Stat3^+/+^* and *Stat3^-/-^* ESCs incubated either with DMSO or CoCl2. H: PLA of STAT3 and HIF1α in *Stat3^+/+^* and *Stat3^-/-^* ESCs incubated either with DMSO or CoCl2. Representative pictures (H) and quantification (H’). Scale bar = 200 μm. *p<0.05; **p<0.01; ***p<0.001; ****p<0.0001; ns = not significant.

**Fig. S3:**
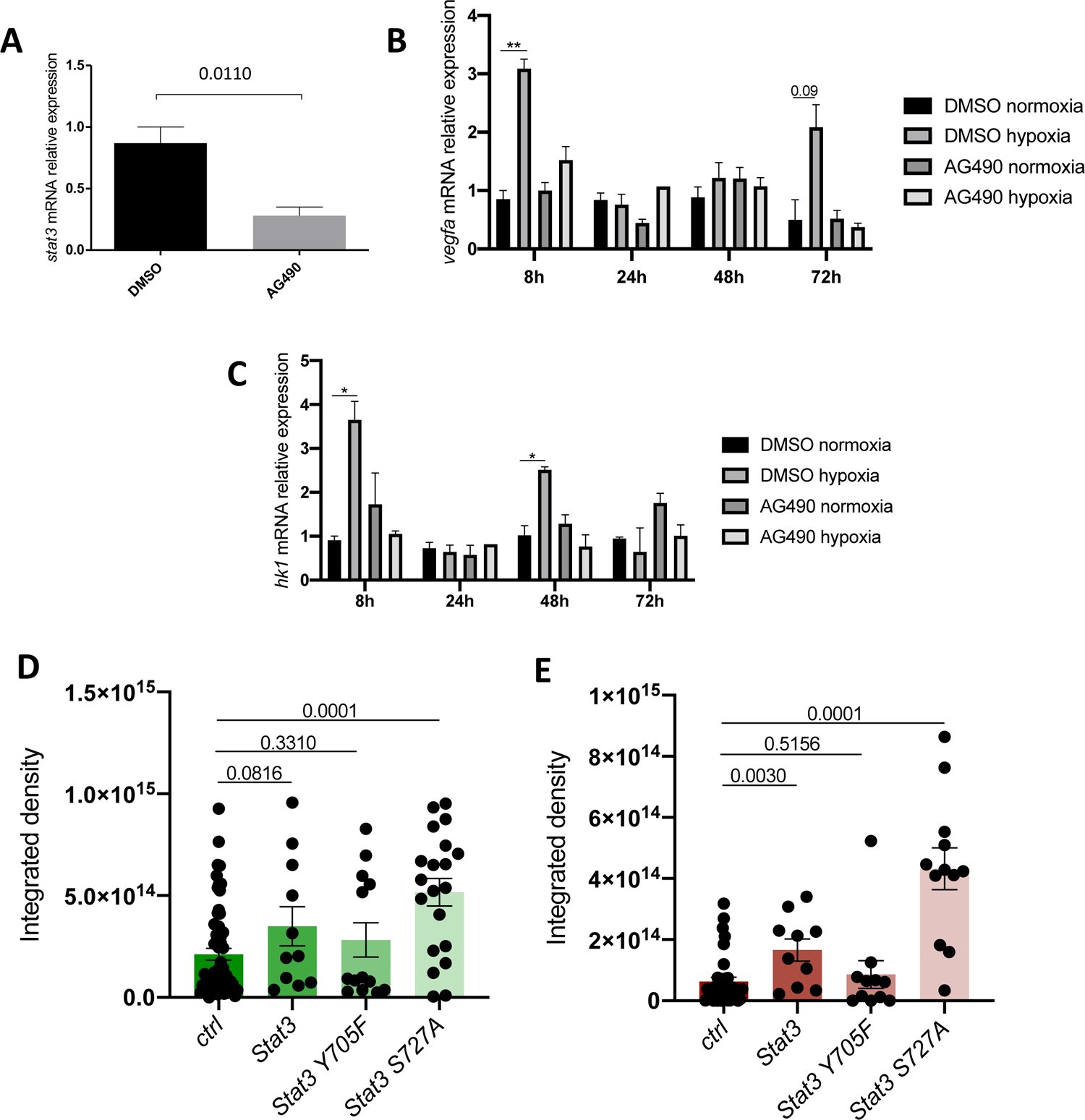
AG490 blocks transcriptional response to hypoxia. A: *stat3* expression level of 6-dpf larvae treated with either DMSO and 50 μM AG490 from 3 dpf to 6 dpf. B: *vegfa* expression of larvae incubated in normoxia or hypoxia and treated with either DMSO and 50 μM AG490 for 8, 24, 48 and 72 hours. C: *hk1* expression of larvae incubated in normoxia or hypoxia and treated with either DMSO and 50 μM AG490 for 8, 24, 48 and 72 hours. D: EGFP fluorescence quantification of *Tg(7xStat3-Hsv.Ul23:EGFP)^ia28^* reporter larvae injected with control solution, *Stat3, Stat3 Y705F, Stat3 S727A* mRNAs. E: mCherry fluorescence quantification of *HRE:mCHerry* reporter larvae injected with control solution, *Stat3, Stat3 Y705F, Stat3 S727A* mRNAs. Mean±SEM. *p<0.05, **p<0.01.

**Fig. S4:**
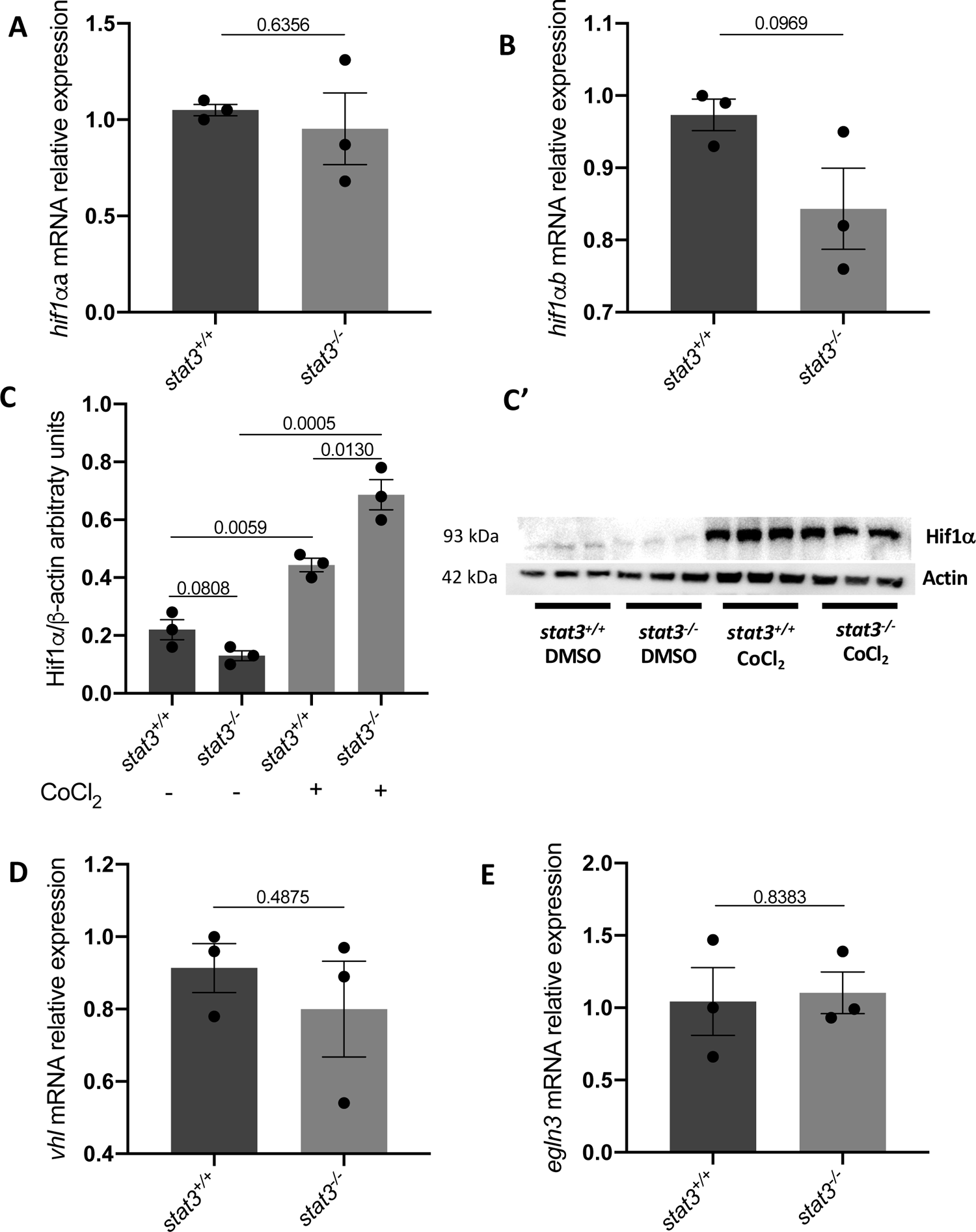
*hif* genes and Hif1α stabilization are not affected in *stat3* knock-out larvae. A: *hif1αa* expression in 6-dpf *stat3^+/+^* larvae compared to *stat3^-/-^* larvae. B: *hif1αb* expression in 6-dpf *stat3^+/+^* larvae compared to *stat3^-/-^* larvae. C: western blot analysis of Hif1α protein in *stat3^+/+^* and *stat3^-/-^* 6-dpf larvae treated either with DMSO or CoCl2. Quantification (C) and representative pictures (C’). D: *vhl* expression in 6-dpf *stat3^+/+^* larvae compared to *stat3^-/-^* larvae. E: *phd3* expression in 6-dpf *stat3^+/+^* larvae compared to *stat3^-/-^* larvae. Mean±SEM. **p<0.01, ***p<0.001, ns=not significant.

**Fig. S5:**
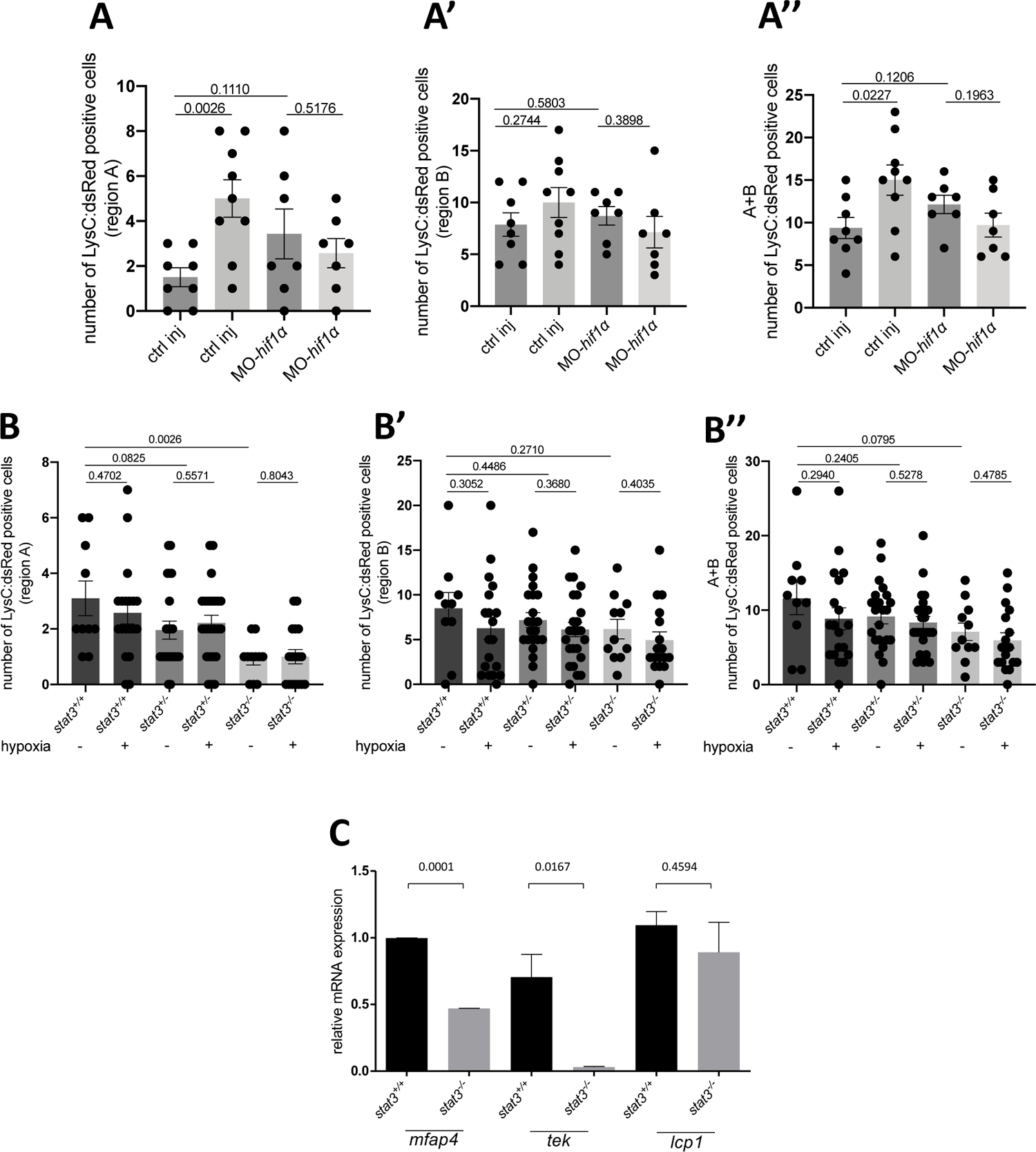
Analysis of macrophage mobilization in *stat3* mutants. A: count of macrophages in region A (A), in region B (A’) and in total (A’’) in ctrl and MO-*hif1a* injected larvae after incubation either in normoxia and in hypoxia. B: count of macrophages in region A (B), in region B (B’) and in total (B’’) in *stat3^+/+^, stat3^+/-^* and *stat3^-/-^* larvae (obtained from the breeding between *stat3^+/-^* parents) incubated either in normoxia and in hypoxia. C: *mfap4, tek* and *lcp1* expression in *stat3^+/+^* and *stat3^-/-^* 6-dpf larvae. *p<0.05; **p<0.01; ***p<0.001; ns = not significant.

## Notes

### Competing Interest Statement

The authors have declared no competing interest.

